# “Shin in the eye”: Au-CS@FNDs as contact lens additives for blocking UV irradiation, bacterial keratitis, and corneal neovascularization therapies

**DOI:** 10.1101/2023.04.11.536356

**Authors:** Linyan Nie, Xiaowen Hu, Yunxiao Zhao, Fan Wu, Yaran Wang, Lei Chen, Yi Wang, Yong Liu

## Abstract

Ophthalmic disease treatment remains a significant problem globally, resulting in poor vision and blindness. UV irradiation and bacterial infection may cause severe damage to the corneal, leading to vision loss within a few days. Corneal neovascular abnormally growth that bock lights reach to eyes, causes low vision. Thus it requires urgent and efficient clinical treatment. Contact lenses play an essential role in treating ophthalmic issues. In this work, we synthesized the chitosan-stabilized Au nanoparticles with a simple method. The Au nanoparticles were further physically adsorbed with negatively charged fluorescent nanodiamonds, yielding Au-CS@FNDs. These Au-CS@FNDs particles were proven with excellent UV adsorption, antibacterial properties, and photothermal conversation ability. Furthermore, we embedded Au-CS@FNDs particles into contact lenses to prevent corneal damage from UV light and bacterial infection. Moreover, the Au-CS@FNDs embedded contact lenses were used to inhibit the neovascularization in the Human Vascular Endothelial Cells via the photodynamic effect of Au nanoparticles. To the best of our knowledge, it is the first time that gold and diamond nanoparticles were used as additives in contact lenses, aiming at clinically corneal neovascular. Our results suggest that the controllable photothermal effect of Au-CS@FNDs embedded contact lenses may provide a unique way to intervene the neovascular-induced vision loss.

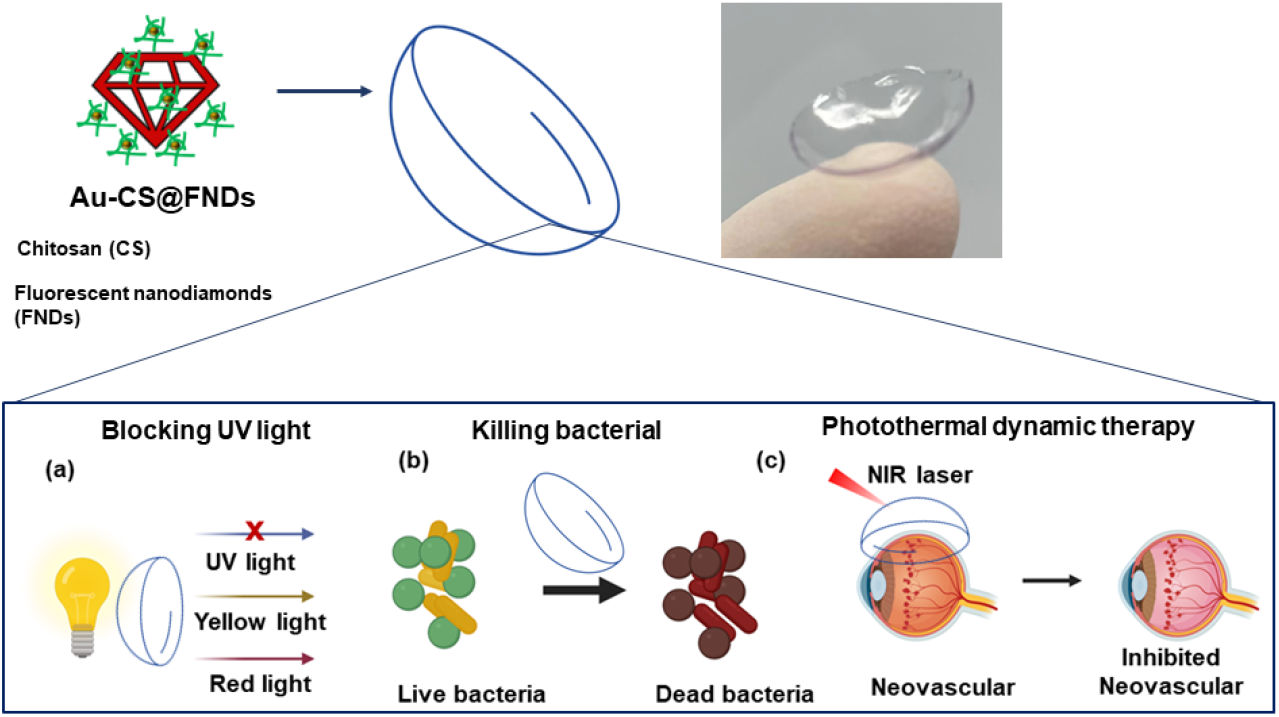

## 1. INTRODUCTION

The eyes are the essential sense organ of human beings. They are related to many vital diseases leading to vision loss or blindness, such as cataracts, glaucoma, age-related macular degeneration, etc. Low vision and blindness significantly impact universally and thus require great attention; these complications can result from many factors, including UV radiation, bacterial infection, abnormal vascular growth in the eye, trauma, nutrition-related diseases ^1, 2^, etc. Eyes are often exposed to an open environment, making them vulnerable to ultraviolet rays and infections. There are increasing reports of UV radiation and intense blue light-induced retina damage, photokeratitis, photophobia, and many other complications ^3, 4^. Besides, bacterial keratitis remains a significant problem worldwide. Infectious keratitis may cause corneal perforation and even can cause severe vision impairment. Multidrugresistant (MDR) bacteria that are not sensitive to broad-spectrum antibiotics result in bacteria keratitis, these bacteria including Staphylococcus aureus (S. aureus), Escherichia coli (E. coli), Pseudomonas aeruginosa (P. aeruginosa), and Streptococcus pneumonia ^5–8^. Among these bacteria, S. aureus is dominant in inducing bacterial infection and requires efficient and quick elimination ^9^. Depending on the size and location of the bacterial infiltration into the corneal epithelium, patients with bacterial keratitis may suffer differently from acute ocular pain, photophobia, conjunctival injection, and vision loss ^10^. Along the same line, Corneal neovascularization (CNV) is another common complication that leads to vision impairment: abnormal blood vessels in the usually clear cornea. The leading underlying cause of CNV is lacking oxygen in the cornea ^11^. Wearing contact lenses for a long time can be attributed to corneal hypoxia ^11,12^. Infections, trauma, and chemical burns can also result in corneal neovascularization.

Contact lenses (CLs) were initially designed for vision correction; with the development of technologies, they are developed into more advanced medical devices. In recent decades, CLs have been intensely researched and developed into various applications ^13–15^, for example, controlled drug delivery for glaucoma ^16–18^, sensing biomarkers ^18^ for cancer diagnoses ^19,20^, and monitoring glucose for diabetes ^21–24^, etc. Nanoparticles like gold ^25^, silver ^25,26^, or silicon ^27^ have been added to contact lenses as additives to make CLs able to achieve more functions, such as dynamic photothermal therapy for posterior capsule opacification (PCO), bacterial killing ^27^, etc. Nanodiamonds are new-class carbon nanomaterials. Due to their unique properties, nanodiamonds have been utilized in various applications, including drug delivery ^28^, biomarkers ^29, 30^, implants ^31^, diagnosis sensors ^32, 33^, and quantum sensing ^34–36^, etc. Nanodiamonds hold high UV absorbance due to the presence of the SP2 structure ^37^. Recently, researchers have been attracted by their excellent UV absorbance ability and utilized diamond particles for UV protection and antioxidant sunscreen products ^38^. Diamonds have been embedded in contact lenses and achieved lysozyme-triggered timolol maleate release for glaucoma treatment ^39^. Fluorescent nanodiamonds (FNDs) are colorless, making them suitable materials as contact lens additives for blocking UV lights. Taking advantage of unique electrical and optical properties, gold nanoparticles are widely used in biomedical applications like imaging, sensing, and targeted drug delivery. The surface plasmon resonance (SPR) of gold nanoparticles enables them to absorb and scatter light ^40–42^; they are also utilized in photodynamic therapy for cancer ^43–46^, bacterial killing ^47–50^, etc. For example, A. E. Salih et al. applied gold-embedded contact lenses to help patients suffering from red-green color vision deficiency, as gold nanoparticles can adsorb green light. L. Dong et al. ^51^ integrated gold nanoparticles in the intraocular lens to treat PCO; irradiated by laser near-infrared (NIR), to eliminate the residual lens epithelial cells, which are the leading cause of PCO. Chitosan (CS), a cationic polymer with permeation-enhancing properties, anticancer, antibacterial, and antioxidant abilities, has been employed in ophthalmic applications ^52^, including sustained drug release ^53^, corneal wound healing, eye drops, ointments, etc.

Aiming at the above three problems (UV irradiation, bacterial infection, and corneal neovascularization), in this work, Figure 1, a biocompatible method was used to synthesize Au nanoparticles, where Au nanoparticles were reduced and stabilized by CS, generating Au-CS nanoparticles ^54, 55^. These positively charged Au-CS nanoparticles further adsorb the negatively charged FNDs, yielding the Au-CS@FNDs nanoparticles that are further embedded in hydroxy ethyl methacrylate (HEMA) based contact lenses. The Au-CS@FNDs implanted contact lenses exhibit excellent UV blocking, in vitro antibacterial, and inhibition of cellular-based neovascular abilities, suggesting Au-CS@FNDs embedded contact lenses are promising for multifunctional ophthalmic diseases.

**Figure 1.**
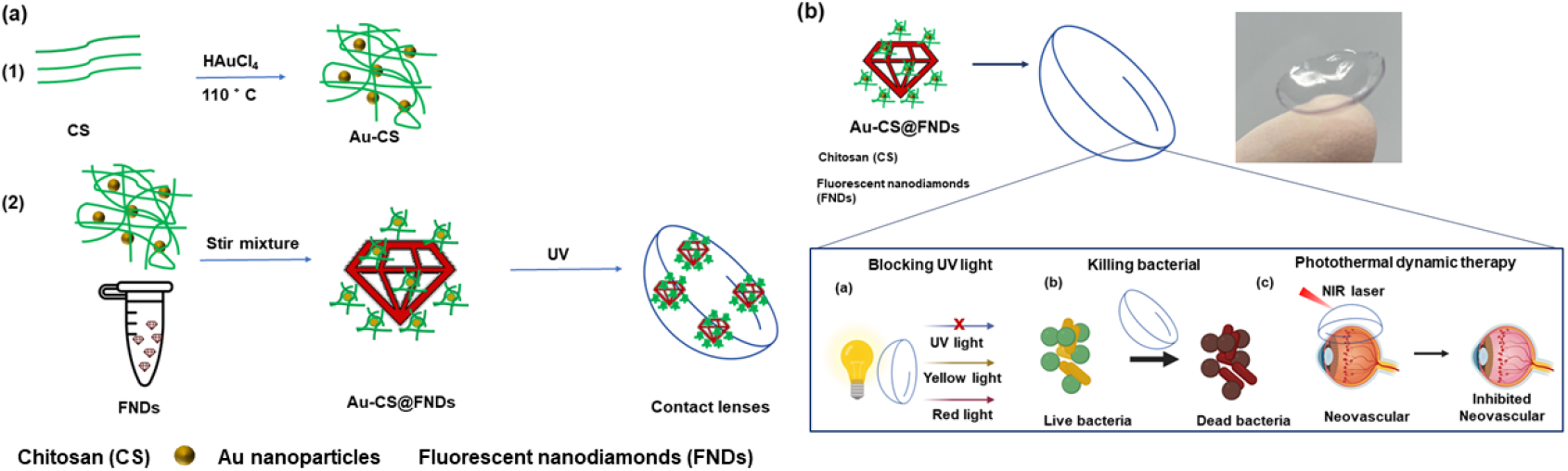
(a) Synthesis of Au-CS@FNDs, and Au-CS@FNDs CLs. (b) schematic illustration of the functions of Au-CS@FNDs CLs.

## 2. RESULTS AND DISCUSSION

### 2.1 Characterization of FNDs, Au-CS, and Au-CS@FNDs nanoparticles

The hydrodynamic size distributions of FNDs, Au-CS, and Au-CS@FNDs were measured by dynamic light scattering (DLS), shown in Figure 2 (a). The hydrodynamic diameter of FNDs, Au-CS, and Au-CS@FNDs is 189 nm (polydispersity index (PDI) is 0.172), 216 nm (PDI = 0.563), and 371.9 nm (PDI = 0.338) respectively. The surface charge of FNDs, Au-CS, and Au-CS@FNDs particles was measured by Zeta potential. In Figure 2. (b), FNDs exhibit a negative surface charge (−9.23 ± 0.036 mV) due to the nitrogen-vacancy centers inside nanodiamonds; Au-CS nanoparticles show a positive surface charge (8.44 ± 0.64) due to the reduction of CS, and the positive surface charge (7.08 ± 0.44 mV) of Au-CS@FNDs suggests that Au-CS were adsorbed on the surface of diamond nanoparticles. Figure 2. (c), the absorption peak of localized surface plasmon resonance (LSPR) of Au-CS@FNDs became broader compared with Au-CS nanoparticles. The adsorption of FNDs, Au-CS, and Au-CS@FNDs (Fig. 2. d) indicates that FNDs and Au-CS@FNDs particles can block light with a short wavelength range (200 - 500 nm), including UV and blue light. The morphology of FNDs, Au-CS, and Au-CS@FNDs is examined by both TEM (Figure 2. (e), (f), (g)) and SEM (Figure S1. (a), (b), (c)). TEM Figure 2. (e) and SEM Figure S1. (a) images indicated that FNDs particles have a flake-like morphology, and the size of FNDs particles is around 100 nm. From TEM (Figure 2. (f)) and SEM (Figure S1. (b)) images, the spherical shape of Au-CS particles was observed. However, the size of Au-CS from EM images indicates Au-CS particles are around 21 nm. This doesn’t agree with the DLS measurement, suggesting that the Au particles have stabilized with a layer of chitosan molecules about 200 nm thick, which expanded in the aqueous phase and shrunk after being dried; this agrees with the previous study ^54, 55^. Figure 2. (e) and Figure S1. (c), the TEM and SEM images both suggest that the Au-CS particles are attached to the flake-like FNDs particles, and the size of Au-CS@FNDs is consistent with that measured by DLS (Figure 2. (b)). The elements containing (Au, oxygen, and nitrogen) of EDS analysis (Figure S2.) suggest the successful synthesis of Au-CS@FNDs particles.

**Figure 2.**
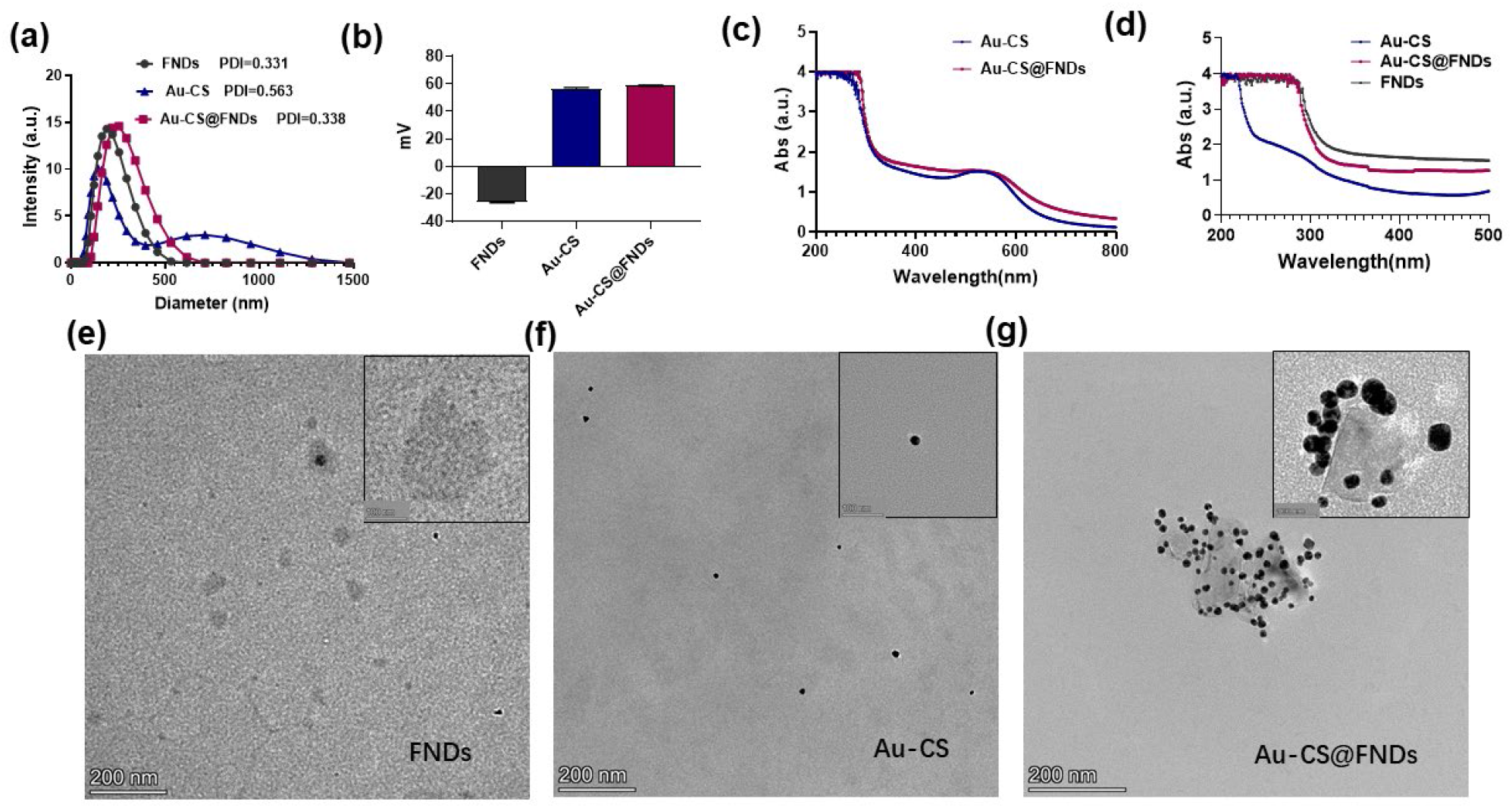
Characterization of FNDs, Au-CS, Au-CS@FNDs nanoparticles. (a) Size distribution of FNDs, Au-CS, Au-CS@FNDs nanoparticles. (b) Surface charges of FNDs, Au-CS, Au-CS@FNDs nanoparticles. (c) and (d) Absorption spectra of FNDs, Au-CS, Au-CS@FNDs nanoparticles. (e), (f), (g) TEM images of FNDs, Au-CS, and Au-CS@FNDs nanoparticles, respectively. The insets are zoomed-in images; scale bars are 100 nm.

### 2.2 Antibacterial activities of FNDs, Au-CS, and Au-CS@FNDs nanoparticles

The contact lens comprises cationic polymer chitosan (CS) and Au nanoparticles. CS and Au nanoparticles were reported to have outstanding antibacterial ability ^56, 57^. The antibacterial ability of CS can attribute to the positive charge provided by the -NH3+ groups, which can interact with the negatively charged cell membranes of a considerable number of both Gram-positive and Gram-negative bacteria ^58^. Au nanoparticles have been used in various antibacterial applications and even used in medical devices to combat multi-drug-resistant bacteria ^59, 60^. Gold nanoparticles do not cause drug resistance compared to antibiotics, as they kill bacteria by altering bacterial membrane potential and inhibiting ATP synthesis. In this work, the antibacterial ability of FNDs, Au-CS, and Au-CS@FNDs nanoparticles was evaluated. *S. aureus* is the dominant pathogenic bacteria that induce bacterial keratitis; they were used to test the antibacterial ability of Au-CS@FNDs. The Gram-negative (*E. coli*) bacteria was also examined as a control. Figure 3. (a) and (b), the quantitative analysis of the bacterial killing ability FNDs, Au-CS, and Au-CS@FNDs nanoparticles, PBS was used as a negative control. Compared with PBS, FNDs didn’t show any differences in the growth of the Gramnegative (*E. coliXen14*) and Gram-positive (*S. aureusXen36*) bacteria strains. Au-CS and Au-CS@FNDs nanoparticles show excellent antibacterial ability against Gram-positive and Gram-negative bacteria strains, indicating their broad spectrum of bacterial killing ability. The SEM images (Figure 3. (c) and (d)) are consistent with the quantitative analysis. In the presence of FNDs, the morphology of *Xen14* and *Xen36* bacteria remained intact. When *Xen14* and *Xen36* bacteria were treated with Au-CS or Au-CS@FNDs nanoparticles, the deformation of these bacteria was observed, suggesting that Au-CS or Au-CS@FNDs nanoparticles damage these bacteria. The growth of Xen14 and Xen36 bacteria on the LB agar plate (Figure S3.) also proved the inhibition ability of Au-CS and Au-CS@FNDs on *Xen14* and *Xen36* bacteria.

**Figure 3.**
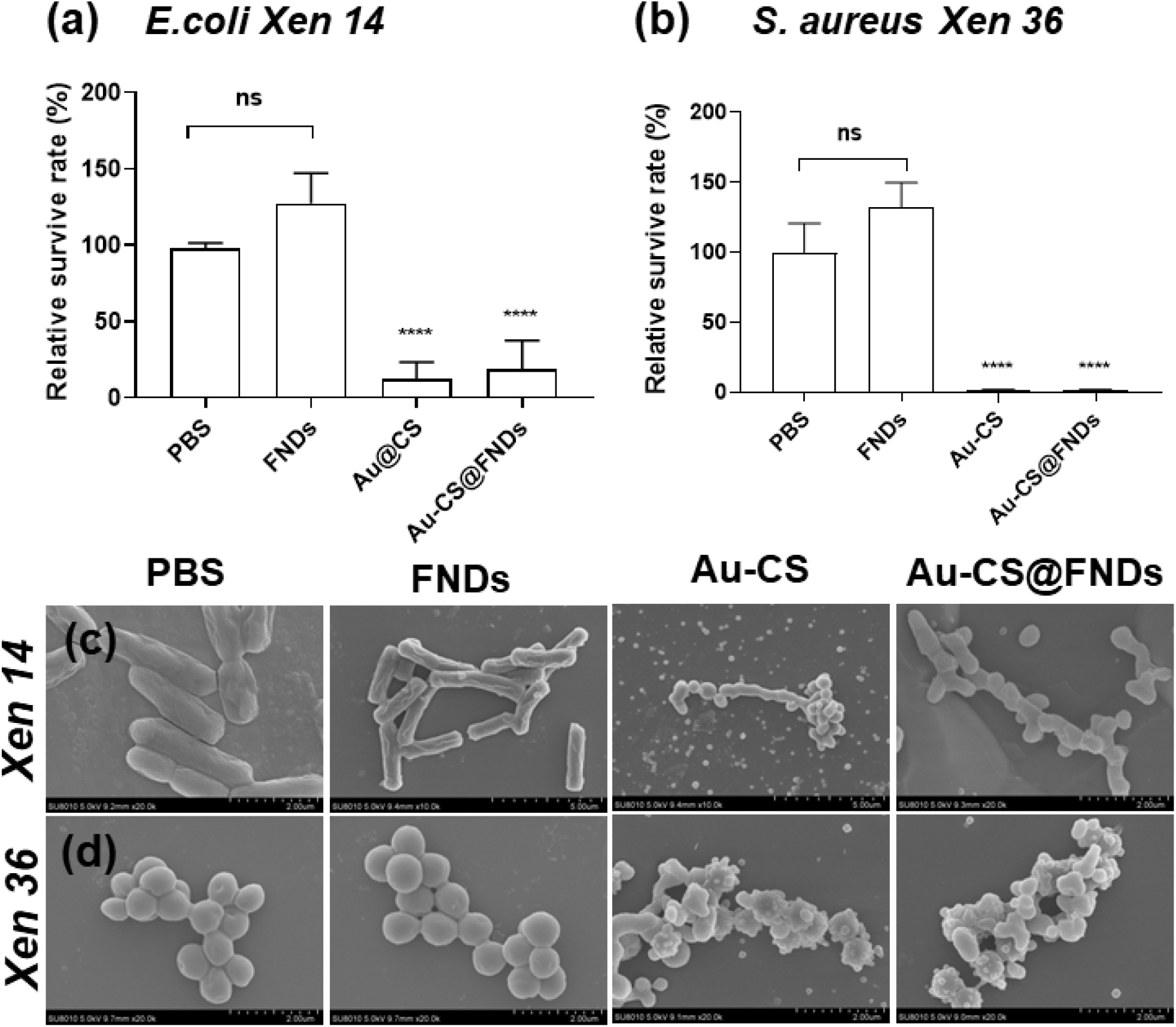
Analysis of the antibacterial ability of FNDs, Au-CS, and Au-CS@FNDs particles. PBS was used as a negative control. An identical amount of FNDs, Au-CS, and Au-CS@FNDs particles were used as in the contact lenses. PBS was used as a negative control. This experiment was repeated three to four times independently. Error bars represent stand deviations. (*** p < 0.0001).

### 2.3 Characterization of Au-CS@FNDs loaded contact lenses

Different volumetric concentrations (2.5%, 5%, 10%, 15%) of Au-CS@FNDs particles were added to the contact lens and investigated by the transmission spectrum (Figure S4.). The results show that Au-CS@FNDs embedded CLs (2.5%, 5%, 10%, 15%) exhibit sound transmission at the wavelength of visible light and remarkable ability to block UV light; 15% of Au-CS@FNDs embedded CLs can also block the blue light (450 – 500 nm) compared to other volumetric concentrations. Besides that, the photodynamic conversation effect depends on the concentration of Au nanoparticles. Next, we examined the reactive oxygen species (ROS) production when HLEC or HCE-2 cells were exposed to Au-CS@FNDs nanoparticles. Excessive ROS will stress the corneal and induce inflammation ^61, 62^, resulting in age-related macular degeneration. Figure S5. compared to the initial group (cells without any treatments), when HLEC or HCE-2 cells were exposed to 15% of Au-CS@FNDs nanoparticles, cellular ROS was significantly decreased, lower amount (2.5%, 5%, or 10%) of Au-CS@FNDs nanoparticles didn’t influence the cellular ROS production. Therefore, to achieve efficient photodynamic therapy by Au-CS@FNDs CLs, 15% volumetric concentration was chosen as the concentration of Au-CS@FNDs particles as CLs additive in this work.

The morphology of pristine and Au-CS@FNDs embedded CLs was evaluated by SEM, shown in Figure 4. (a). The aggregates of Au-CS@FNDs particles were observed in the CLs. A. E. Salih et al. ^28^ also observed the same scenario when embedding gold nanocomposite into a contact lens. Au-CS@FNDs CLs were found to possess more water content than the pristine ones; this could be attributed to chitosan’s strong water absorption moisturizing ability ^64^. Gold nanoparticles have been widely used in photodynamic therapy due to their excellent photothermal conversation effect ^65^. The photothermal conversation effect of FNDs, Au-CS, and Au-CS@FNDs was evaluated (Figure S6.) under different laser power (2 W/cm^2^, 3 W/cm^2^, 4 W/cm^2^, 5 W/cm^2^). The data indicate an excellent photothermal conversation effect of Au-CS and Au-CS@FNDs nanoparticles under NIR laser irradiation at the laser power of 4 W/cm^2^ and 5 W/cm^2^. To avoid overheating, the laser power was chosen as 4 W/cm^2^ for further experiments. Some studies used this laser power to perform photodynamic therapy for curing PCO in the rabbit model ^51^. The photothermal effect of Au-CS@FNDs CLs (Figure 4. (d)) was also tested. The pristine and Au-CS@FNDs embedded CLs were placed in 1 mL PBS for the measurement. Under the 808 nm NIR irradiation (4 W/cm^2^), in the presence of Au-CS@FNDs CLs, the temperature of PBS increased 2. 2 °C within 15 min. Under the same experimental conditions, we didn’t observe any temperature changes in the presence of pristine CLs or PBS alone.

**Figure 4.**
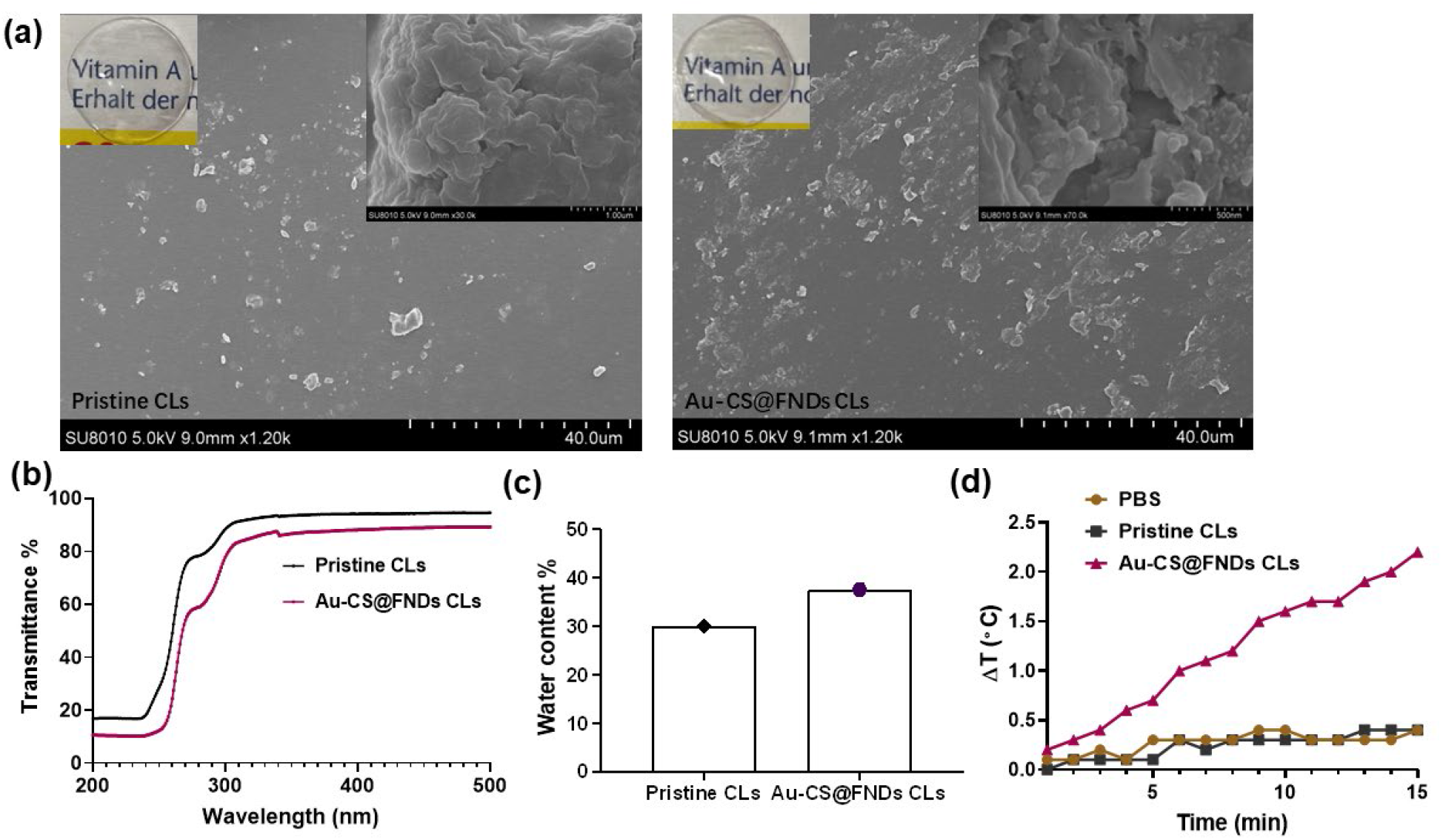
Characterization of Au-CS@FNDs CLs. (a) SEM images of pristine CLs and Au-CS@FNDs CLs. The insets are zoomedin images. (b) Transmission spectra of pristine contact lens and Au-CS@FNDs embedded CLs. (c) The water content of pristine contact lens and Au-CS@FNDs CLs. (d) The photothermal conversion property of pristine CLs and Au-CS@FNDs CLs in PBS under NIR irradiation for 15 min at the laser power of 4 W/cm^2^.

Next, the biocompatibility of FNDs, Au-CS, and Au-CS@FNDs nanoparticles was tested with a CCK8 assay. The concentration of FNDs, Au-CS, and Au-CS@FNDs nanoparticles is identical to the Au-CS@FNDs embedded CLs. The human corneal epithelial cells (HLECs, HCE-2 Cells) were used in this experiment, the cells were incubated with FNDs, Au-CS, and Au-CS@FNDs nanoparticles for one day, three days, and five days; the amount of FNDs, Au-CS, Au-CS@FNDs nanoparticles were identical to what we used for contact additives. 5% of DMSO has used a positive control (PC), leading to cell death. As shown in Figure S7, no differences were observed between the negative control (NC) group and cells exposed to FNDs, Au-CS, and Au-CS@FNDs even after 72 h. The cell viability test suggests excellent biocompatibility of FNDs, Au-CS, and Au-CS@FNDs nanoparticles with HLECs and HCE-2 cells.

### 2.4 Inhibition of HVECs by photothermal effect of Au-CS@FNDs CLs

The growth of new blood vessels from the capillaries and venules of the peripheral vascular plexus can block lights entering the eyes and affect vision ^65, 65^. The corneal neovascular (CNW) can be secondary to excessive reactive oxygen species with eye drops, inflammation, chemistry burns, trauma, etc. CNW is traditionally treated with eye drops; however, the therapeutic efficiency is low due to rapid elimination. Nowadays, nanomaterials with photothermal effects are injected into rabbit corneal to achieve CNW inhibition ^66–69–8^. However, this requires an experienced ophthalmologist. Inspired by the photodynamic therapy on tumor inhibition, the Au-CS@FNDs CLs are employed to inhibit CNW. The human vascular endothelial cells (HVECs) have been used as a model to study vascular inflammation in diabetes ^69^ and CNW ^7^0,71 in this work, we use HVECs to evaluate the inhibition of neovascular by the photothermal effect of Au-CS@FNDs CLs. For the in vitro experiment, HVECs were seeded in a 24-well plate at a 90% confluence; after they were attached to the bottom of the plate for kept in the incubator overnight, pristine CLs and Au-CS@FNDs CLs were carefully placed on the top of the cells and irradiated with an 808 nm laser (4 W/cm^2^) for 10 min, followed by staining with Calcein-AM to observe the viable cells, Calcein-Am stains live cells with green fluorescence. In Fig.5., in the presence of Au-CS@FNDs CLs, there are a few cell residues that remain green signal, indicating photothermal-killing effects on cells from the Au-CS@FNDs CLs. Under the same irritation condition, the NIR laser didn’t show any killing on cells or cells in the presence of pristine CLs.

**Figure 5.**
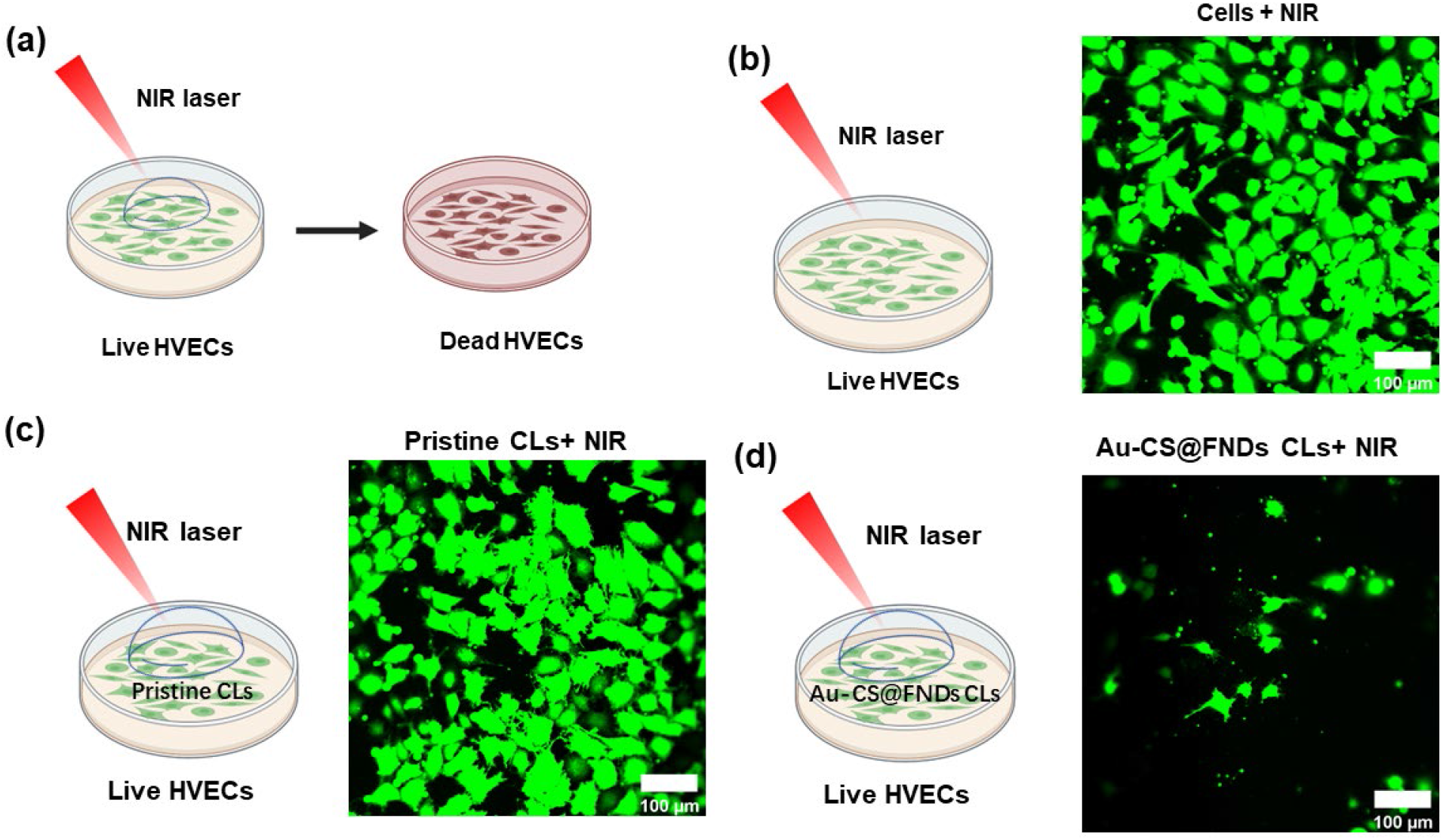
The inhibition ability of Au-CS@FNDs CLs against HVEC cells. (a) schematic illustration of Au-CS@FNDs embedded contact lens-based photodynamic effect on the inhibition of HVECs. (b) (c) (d) control, pristine CLs, Au-CS@FNDs CLs under NIR laser irradiation for 10 min at the laser power of 4 W/cm^2^. Green: Calcein-AM for live cell staining, scale bars are 100 μm. This experiment was repeated three times independently.

## 3 CONCLUSIONS

Contact lenses are smart medical devices beyond their primary functions; they are involved in various applications in ocular diseases. For the first time, we fabricated contact lenses with Au-CS@FNDs additives that possess multifunction: anti UV irradiation, antibacterial infection, and inhibit corneal neovascular through photodynamic therapy. The great antibacterial activities of Au-CS@FNDs particles can prevent bacterial corneal keratitis caused by wearing contact lenses. The photothermal effect of Au-CS@FNDs CLs on killing HUVE cells makes the Au-CS@FNDs CLs promising for noninvasive therapy of corneal neovascular. Taken together, contact lenses with Au-CS@FNDs additives are not only suitable for daily wear that protects eyes from irritation, but they also promise therapeutic devices.

## 4 EXPERIMENTAL METHODS

### 4.1 Materials

Fluorescent nanodiamonds (FNDs) with a hydrodynamic diameter of 100nm were purchased from Adamas Nanotechnologies, Brownleigh Drive Raleigh, NC, USA. The diamonds were produced by grinding at high-pressure high temperature, and their surface is oxygen-terminated due to an acid cleaning treatment with oxidizing acids by the manufacturer. Chitosan (MW=50,000), hydrochloroauric acid solution (HAuCl4), 2-hydroxyethyl methacrylate (HEMA, 99%), ethylene glycol dimethacrylate (EGDMA, 98%), and 2-hydroxy-2-methyl-1-phenyl-1-propanone (HMPP, 97%) were all purchased from Shanghai McLean Biochemical Technology Co., Ltd. Calcein /PI Live/Dead Viability/Cytotoxicity Assay Kit) from Beyotime, Shanghai, China. DCFDA / H2DCFDA - Cellular ROS kit and Cell Counting Kit 8 (WST-8 / CCK8) (ab228554) were purchased from Abcam (Shanghai, China).

### 4.2 Preparation of Au-CS@FNDs nanoparticles

The synthesis of Au-CS nanoparticles was described previously ^54, 55^ with slight modification. Briefly, 20 mL of 1.8 % chitosan solution dissolved in acetic acid was heated under reflux until boiling. Then 10 mM HAuCl4 solution was added with vigorous stirring. The collected Au-CS was dispersed in deionized water. Then Au-CS nanoparticles and FNDs were stirred and mixed for 1 h at 4°C in a refrigerator, allowing positively charged Au-CS nanoparticles to physically adsorb negatively charged FNDs particles.

### 4.3 Characterizations of nanoparticles

A Malvern ZetaSizer Nnaosystem (Malvern Instruments Ltd., Malvern, UK) was used to evaluate the hydrodynamic size distribution and surface charge of FNDs, Au-CS, and Au-CS@FNDs nanoparticles. The absorption spectra were obtained using an ultraviolet spectrophotometer (UV1900i, SHIMADZU, China). Transmission electron microscope (TEM, Talos F200S, Thermo Scientific, China) and energy-dispersive X-ray spectra (EDS) were used to characterize Au-CS@FNDs nanoparticles.

### 4.4 Preparation of Au-CS@FNDs loaded contact lens

The preparation of Au-CS@FNDs loaded contact lens was described earlier ^18^. Briefly, a mixture (1 mL) containing 2-hydroxyethyl methacrylate (HEMA), acrylic acid (AAc) (130 μL), 2,2-diethoxyacetophenone (DEAP), and trimethylolpropane trimethacrylate (TMPTMA), was irradiated under a 365 nm UV lamp (UVATA Technology Co., Ltd) to polymerize the contact lenses. Different concentrations (2.5%, 5%, 10%, 15%) of Au-CS@FNDs were loaded to HEMA based contact lens.

### 4.5 Characterizations of Au-CS@FNDs embed contact lens

The transmittance of control and Au-CS@FNDs loaded contact lenses were analyzed using an ultraviolet spectrophotometer (UV1900i, SHIMADZU, China). The transmittance of the contact lenses was measured in the wavelength range of 200 – 800 nm. The water content analysis of the control and Au-CS@FNDs loaded contact lenses was performed using a vacuum desiccator. The surface morphologies of control and Au-CS@FNDs loaded contact lenses were evaluated using a scanning electron microscope (SEM, Phenom, Phenom Pharos, the Netherlands).

### 4.6 Bacterial culture

*Escherichia coli* (*E. coliXen14,* WS2572), a Gram-negative clinical bacterium isolates from Weihenstephan Culture Collection, and *Staphylococcus aureus* (*S. aureusXen36*), a Gram-positive parental strain from the American Type Culture Collection (ATCC) were purchased from PerkinElmer Inc., Waltham, MA, USA, and were grown in Luria-Bertani (LB) broth medium. Bacterial cultures of *E. coli* and S. *aureus* were prepared by suspending a colony in a 10 mL LB medium and incubated at 37°C for 24 h aerobically. Subsequently, 2.5 mL were transferred into 50 mL TSB in a ratio of 1:20 (v / v) followed by 24 h incubation at 37°C. Next, the cultures were washed twice with 10 mL sterile phosphate-buffered saline (PBS), alternated by centrifugation at 10°C and 6500 rpm (Beckman Coulter JLA 16.250). Enumeration of bacteria in the suspension was quantified with a Bürker-Türk counting chamber.

### 4.7 Cell culture

Human corneal epithelial cells (HCE-2 cells) were cultured in Dulbecco’s Modified Eagle Medium/Nutrient Mixture F-12 (DMEM/F-12) medium, supplemented with 10% fetal bovine serum (FBS), 1% penicillin/streptomycin (10,000U/mL penicillin, 10,000 ug/mL streptomycin), 5 μg/mL insulin. Human lens epithelial cells (HLECs) were cultured in DMEM/F-12 medium supplemented with 10% fetal bovine serum (FBS), 1% penicillin/streptomycin (10,000U/mL penicillin, 10,000 ug/mL streptomycin). Human vascular endothelial cells (HUVCs) were cultured in Endothelial cell growth medium-2 (EGM-2) supplemented with Growth Factors, CC-4176, Lonza Inc.). These cells were cultured at 37°C and 5% CO2 in an incubator.

### 4.8 Cellular reactive oxygen species measurement

To investigate the influences of Au-CS@FNDs nanoparticles on human corneal, cellular ROS was examined with a DCFDA / H2DCFDA - Cellular ROS. The measurement is based on the diffusion of DCFDA into the cell; cellular esterases then deacetylate it to a non-fluorescent compound, which ROS later oxidizes into 2’, 7’ -dichlorofluorescein (DCF); DCF is highly fluorescent. HCE-2 cells and HLECs were seeded in a 96-well microplate at a seeding density of 5,000 cells/well in a serum-containing medium. The cells were kept in an incubator overnight, allowing them to attach; the medium was replaced with a 200 uL serum-free medium containing (2.5%, 5%, 10%, 15%) Au-CS@FNDs particles following incubation for 24 h. After incubation, the medium containing nano-particles was removed, and cells were rinsed with PBS and stained with DCFDA. A microplate reader measured the fluorescence with excitation/emission at 485 nm/535 nm (WM2016005, ThermoFisher, the US). Cells that didn’t stain with DCFDA were used as a background substrate, and cells without any treatment and stained with DCFDA were used to measure the initial ROS generated by normal cellular metabolism.

### 4.9 Cell cytotoxicity

HCE-2 cells and HLECs were used to test the in vitro toxicity of the FNDs, Au-CS, and Au-CS@FNDs particles used for the contact lenses. Briefly, HCE-2 cells and HLECs were plated in a 96-well microplate at a seeding density of 5,000 cells/well in a serum-containing medium. The cells were kept in an incubator overnight, allowing them to attach; the medium was replaced with a 200 uL serum-free medium containing FNDs, Au-CS, and Au-CS@FNDs particles following incubation for one day, three days, and five days, respectively. The concentration of FNDs, Au-CS, and Au-CS@FNDs particles is identical to the number used in the contact lenses contacting 15% of Au-CS@FNDs particles. After incubation, 10 uL CCK8 was added to each well, and following incubation for 3 h at 37 °C, the absorbance was recorded at 450 nm using a microplate reader (WM2016005, ThermoFisher, the US). The concentration of FNDs, Au-CS, and Au-CS@FNDs particles are identical as used in contact lenses. Cells without any treatment were used as a negative control, and 5% dimethyl sulfoxide (DMSO) was used as a positive control, as DMSO induces cell death.

### 4.10 In vitro antibacterial property of FNDs, Au-CS, Au-CS@FND nanoparticles

S. aureus and E. coli were used for antibacterial experiments, and the bacterial intensity was adjusted to a proper concentration at 1 x 107 CFU/mL. The samples (FNDs, Au-CS, Au-CS@FNDs) were added to a 5 mL of Luria Bertani broth (LB) containing 5 mL bacterial solution (100 uL) and incubated for 6 h. After that, the bacterial suspensions were collected and diluted in PBS, evenly spread on the agar plate, and incubated for 24 h at 37 °C. Finally, the number of bacterial colonies on the agar was counted. The following formula calculated the bacterial survival rate:

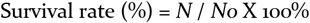

### 4.11 Morphological characterization of bacteria

The bacterial suspensions of *S. aureus* and *E. coli* that were treated with PBS, FNDs, AU-CS, and Au-CS@FNDs nanoparticles as described above 2.10 were collected, and placed on a silicon wafer, kept in the incubator for 30 min allowing them to attach. Subsequently, the samples were fixed with 4% Paraformaldehyde solution and dehydrated with a graded series of ethanol (30%, 50%, 70%, 90%, 100%) for 15 min. the samples were then dehydrated again in a Hitachi Model HCP-2 critical point dryer. Prior to SEM imaging, the samples were coated with gold palladium.

### 4.12 Photothermal conversation effect evaluation

To examine the photothermal conversation of these nanoparticles (FNDs, Au-CS, Au-CS@FNDs), the samples were placed under a NIR laser transmitter (Lasever, Inc., China) and an Infrared thermal imager (FLIR) with different laser power density (2 W/cm2, 3 W/cm2, 4 W/cm2, 5 W/cm2) respectively. Au-CS@FNDs loaded contact lenses were placed in phosphate-buffered saline (PBS) in a 48-well microplate to elevate their photothermal conversation effect.

### 4.13 HVECs inhibition by photothermal effect of Au-CS@FNDs embed contact lens

The experiment method is described by C. Zhang et al. [56] with slight modifications. HVECs were used as a cell model for the in vivo corneal neovascular experiments. HVECs were seeded in a 6-well microplate at a 90% confluence and kept in an insulator, allowing them to attach to the bottom. The medium was replaced with 1mL of fresh serum-free cell culture media, and Au-CS@FNDs loaded contact lenses were placed in the well; three groups were randomized and established: cells with NIR irradiation, pristine CLs with NIR irradiation, Au-CS@FNDs embedded contact lenses with NIR irradiation. The contact lenses were carefully placed into the wells and kept in the incubator for 12 h. Subsequently, these three groups were irradiated with a NIR laser (808 nm, 4 W/cm2) for 10 min. To examine the inhibition of HVECs, a Live/dead staining assay was carried out using a Cal-cein-AM (CM). After staining, the cells were imaged with confocal microscopy (Nikon, A1, Japan). The assay is based on the integrity of the cell membrane. Calcein-AM is membrane permeable; inside live cells, it emits green fluoresce (ex/em = 490/ 515 nm).

### 2.14 Statistical analysis

Prism GraphPad 8 was used to analyze data unless indicated; one-way ANOVA was used to compare data among groups, and a T-test was used to compare two groups. The data were presented as median with interquartile range and * p < 0.05, ** p< 0.001, **** p< 0.0001 were defined as significant differences.

## Supporting information

https://kuleuven-my.sharepoint.com/personal/linyan_nie_kuleuven_be/_layouts/15/onedrive.aspx?login_hint=linyan%2Enie%40kuleuven%2Ebe&id=%2Fpersonal%2F

## ASSOCIATED CONTENT

**Supporting Information**

## AUTHOR INFORMATION

### Corresponding Author

**\*(L. Nie.),**

**\*(Y. Wang.),**

**\*(Y. Liu.).**

### Author Contributions

Linyan Nie designed and conceived the study, acquired and analyzed the data, and drafted the original manuscript. Yunxiao Zhao prepared contact lenses under the supervision of Yi Wang. Xiaowen Hu performed SEM imaging of bacteria and nanoparticles. Fan Wu performed TEM imaging of nanoparticles. Yaran Wang performed SEM imaging of contact lenses. Lei Chen helped Linyan Nie prepare Au-CS nanoparticles. Linyan Nie, Yi Wang, and Yong Liu wrote the article. All authors have read and agreed to the published version of the manuscript.

### Funding Sources

#### Acknowledgment

The authors acknowledge Eye hospital, WMU Zhejiang Eye hospital for the cells of HLEC, HCE-2, and HUVECs. National Natural Science Foundation of China (Grant No. 52003184) and Startup Fund of Wenzhou Institute, University of Chinese Academy of Sciences (Grant No. WIUCASQD2021022) for Yong Liu. Zhejiang Province Natural Science Fund for Distinguished Yong Scholars (LR19H180001) for Yi Wang.

### Notes

The authors declare no competing financial interest.

