## Supplementary material for "“Shin in the eye”: Au-CS@FNDs as contact lens additives for blocking UV irradiation, bacterial keratitis, and corneal neovascularization therapies": https://kuleuven-my.sharepoint.com/personal/linyan_nie_kuleuven_be/_layouts/15/onedrive.aspx?login_hint=linyan%2Enie%40kuleuven%2Ebe&id=%2Fpersonal%2F

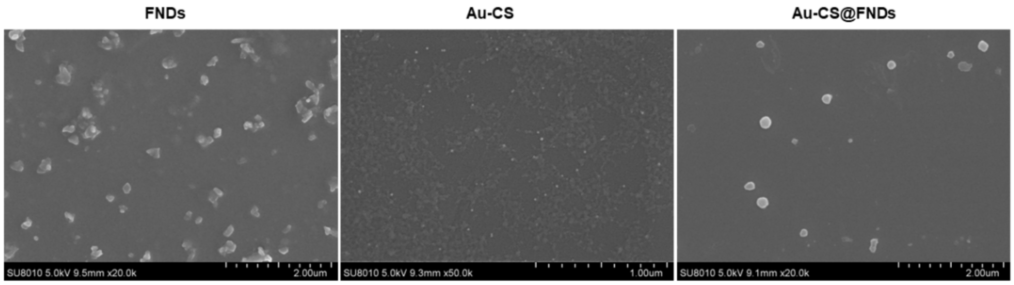


**Figure S1. SEM images of FNDs, Au-CS, and Au-CS@FNDs particles.**

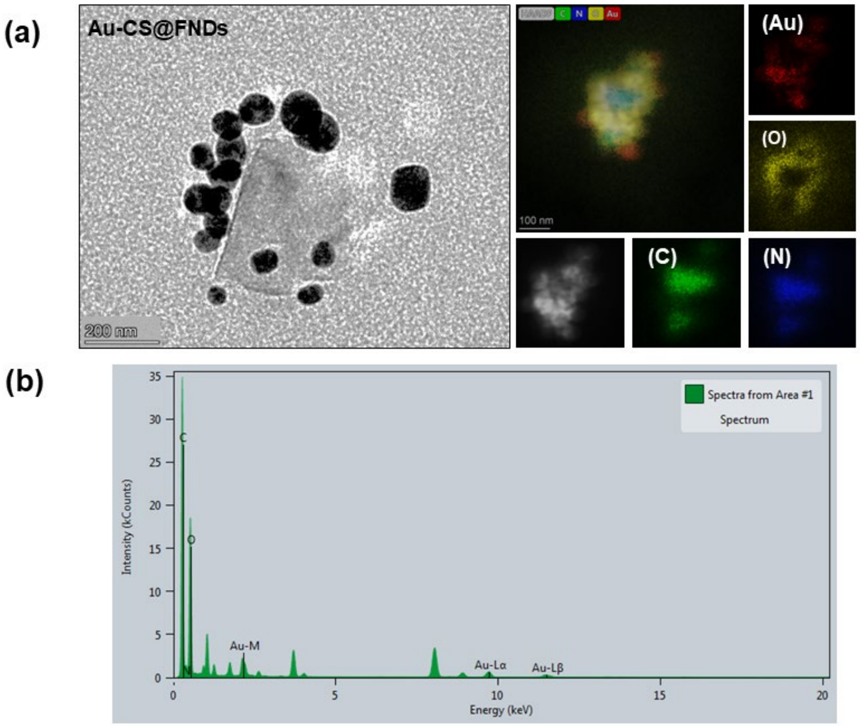


**Figure S2. Morphology and EDS analysis of Au-CS@FNDs nanoparticles. (a) TEM micrograph image of Au-CS@FNDs particles. Red: Au, orange: oxygen, green: carbon, and blue: nitrogen of element mapping of Au-CS@FNDs nanoparticles. (b) quantitively analysis of elements of Au-CS@FNDs nanoparticles.**

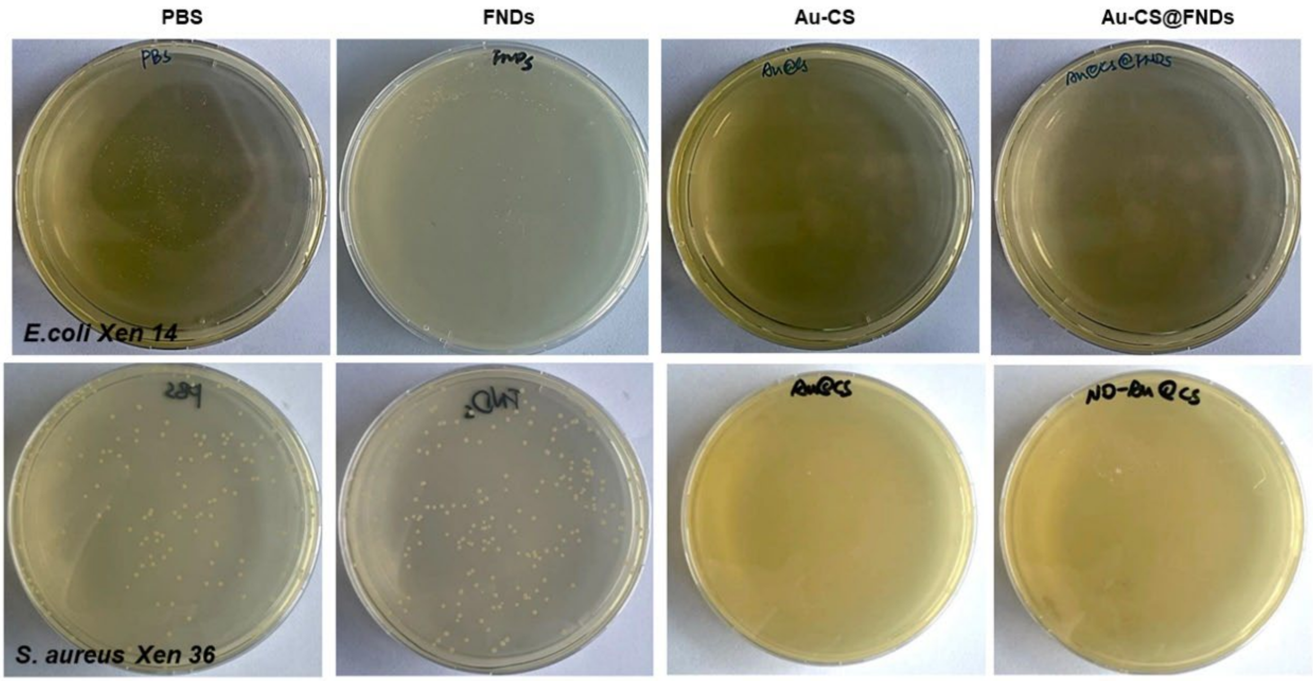


**Figure S3.** Images of FNDs, Au-CS, and Au-CS@FNDs particles killing bacteria; PBS was used as a negative control.

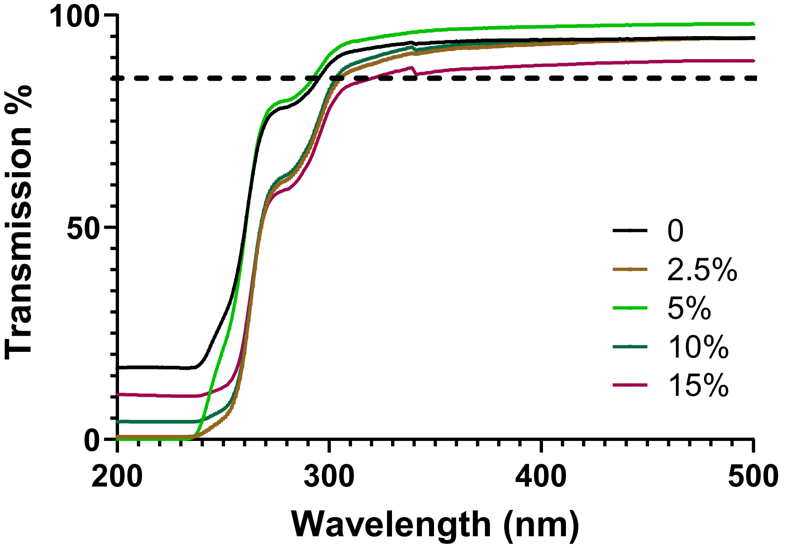


**Figure S4.** The transmission of different volumetric concentrations (2.5%, 5%, 10%, 15%) of Au-CS@FNDs embedded contact lenses.

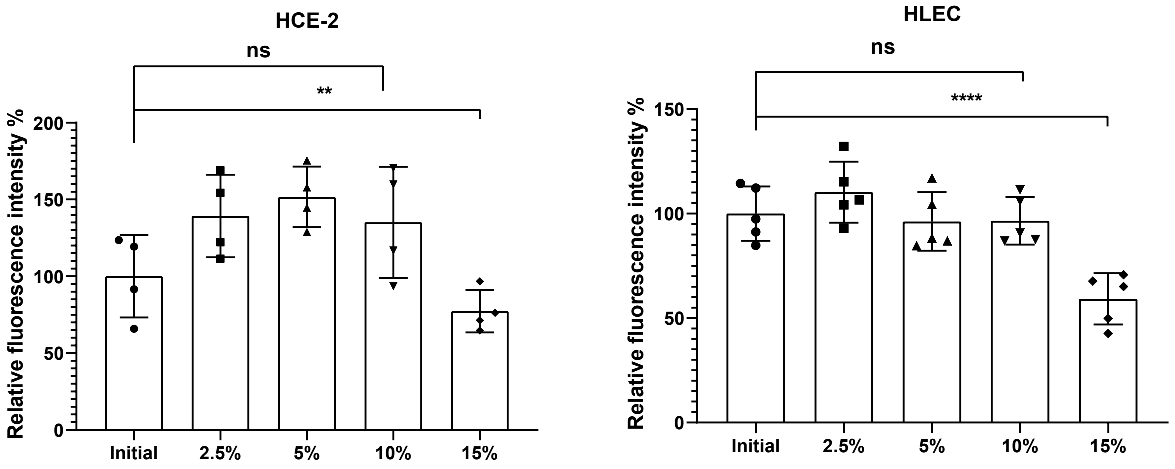


Figure S5. Cell ROS production of Au-CS@FNDs nanoparticles incubated with HCE-2 cells and HLEC cells. HCE-2 cells and HLECs were incubated with FNDs, Au-CS, and Au-CS@FNDs particles for 24 h, respectively. Different volumetric concentrations (2.5%, 5%, 10%, 15%) of Au-CS@FNDs particles were used to incubate with cells. The amount of these particles is identical to that used in the contact lens. Cells without any treatment were used to measure the initial amount of cellular ROS generation, blank wells were used as backgrounds. The experiment was repeated three times independently; error bars represent stand deviations. (**** $p < 0.0001$).

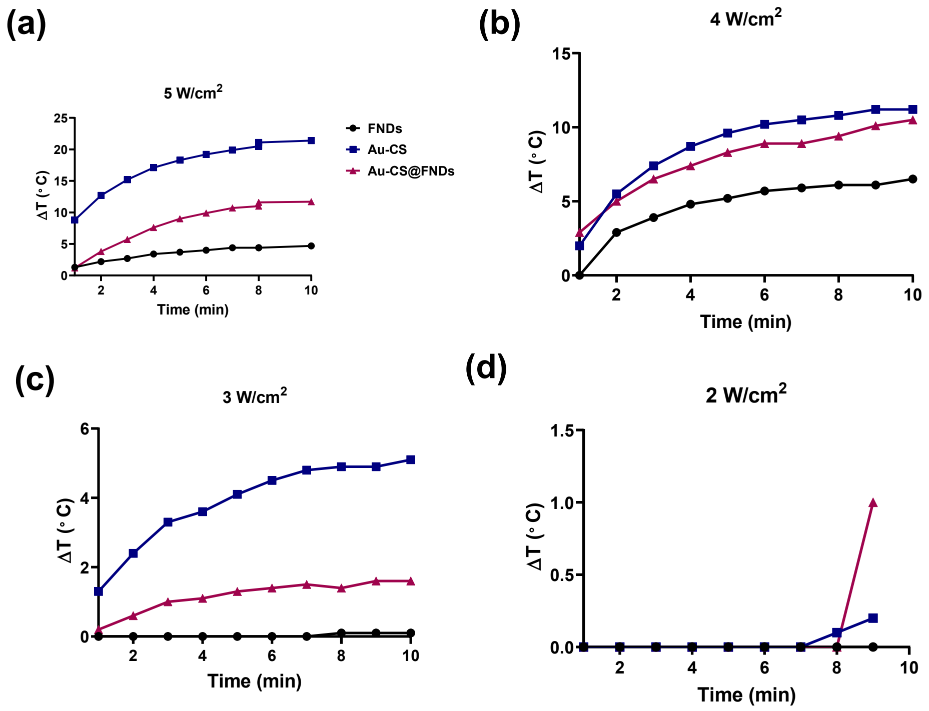


Figure S6. The photothermal effect of FNDs, Au-CS, and Au-CS@FNDs nanoparticles under different laser power for 10 min (NIR: 808 nm). Black: FNDs, blue: Au-CS, rose red: Au-CS@FNDs.

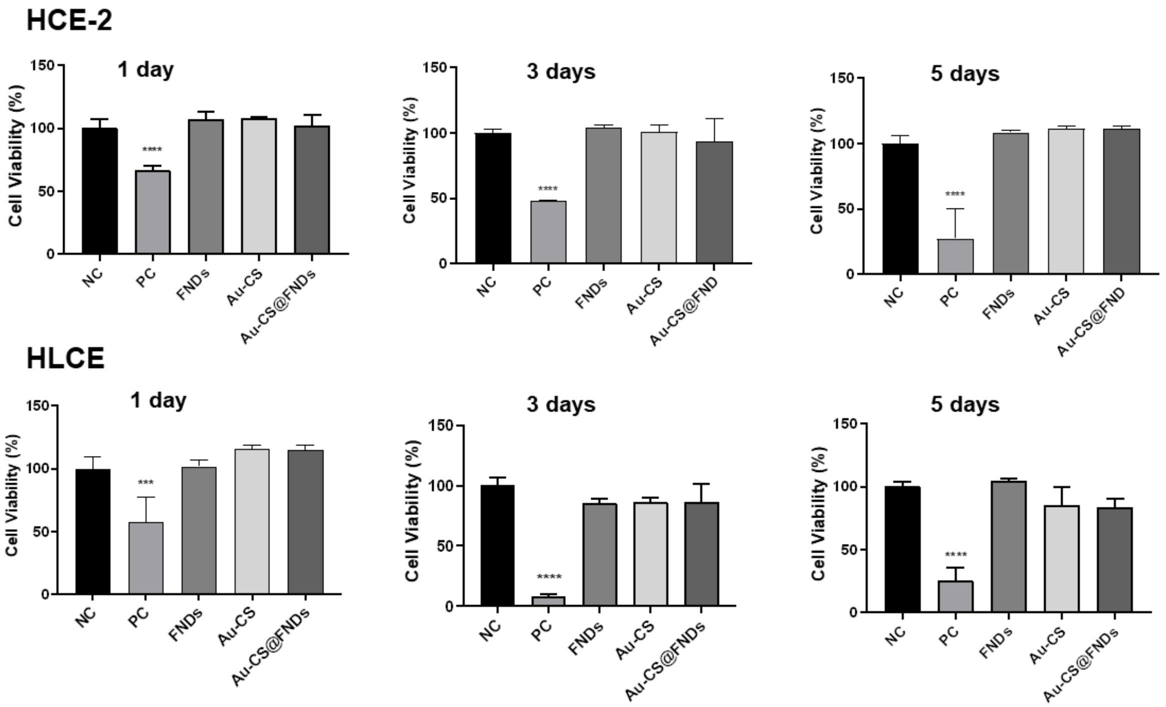


**Figure S7. Cell viability test of FNDs, Au-CS, and Au-CS@FNDs nanoparticles. Cell visibility test of HCE-2 cells and HLECs incubated with FNDs, Au-CS, and Au-CS@FNDs particles for different time points, respectively, the amount of these particles is identical as used in the contact lens. NC: negative control, cells without any treatment; PC: positive control, cells treated with 5% DMSO. This experiment was repeated three times independently. The experiment was repeated three times independently; error bars represent stand deviations. (**** $p < 0.0001$).**
